# GPU acceleration of cell-based simulations in Chaste using FLAME GPU 2

**DOI:** 10.64898/2026.01.13.699201

**Authors:** Matthew Leach, Peter Heywood, Alexander G. Fletcher, Paul Richmond

**Author notes:** **CONTACT INFORMATION** M.L. P.H. A.F. P.R.

## Abstract

Chaste is an open-source C++ library providing a general-purpose framework for cell-based simulations of biological tissues. It has been applied to a wider range of biological processes, including morphogenesis, carcinogenesis, and wound healing. Such simulations often involve numerous mechanical interactions between neighbouring cells, making them computationally demanding. Graphical Processing Units (GPUs), with their highly parallel architectures, offer a powerful means to accelerate these computations, enabling larger, more detailed simulations and improving research productivity.

FLAME GPU 2 is a GPU-accelerated simulator for domain-independent complex systems that maps formal agent descriptions written in scripting language to optimized CUDA code. In this work, FLAME GPU 2 is integrated with Chaste to accelerate force calculations in a class of cell-based simulations, demonstrating the feasibility of GPU acceleration within existing CPU-based frameworks. The GPU-accelerated implementation is validated against the original CPU version, achieving up to 93.6x speedup in force calculations and 3.72x speedup for full simulations across various cell population sizes. Moreover, the approach enables smaller mechanics time steps, without incurring significant data transfer overhead, thereby improving the accuracy of mechanical modelling. This enhancement increases the fidelity of cell position calculations in non-equilibrium simulations and improves dynamic accuracy as cells approach equilibrium.

## INTRODUCTION

Agent-based modelling approaches are powerful tools for understanding and predicting the dynamic behaviour of biological tissues in health and disease. By representing individual cells as discrete entities with defined interaction rules, these models can capture complex multicellular phenomena such as morphogenesis [1], tumour growth [2], and tissue repair [3]. Consequently, they are increasingly used to explore emergent behaviours across spatial and temporal scales that are otherwise difficult to study experimentally.

Numerous cell-based simulators aim to reproduce or predict multicellular behaviour, often using agent-based models that vary in force laws, integration methods, domain discretisation approaches and flexibility. PhysiCell [4] employs an off-lattice approach for 2D and 3D domains, with intracellular behaviour defined by SBML models, Boolean networks, or diffusion flux balance analysis, and intercellular interactions modelled by treating cells as spheres subject to potential forces. Morpheus [5] uses a Cellular Potts Model [6] on 2D and 3D regular lattices, where intracellular behaviours are governed by SBML models [7] or ODEs, and intercellular forces are computed from a Hamiltonian-like functional. BioDynaMo [8] adopts an off-lattice discretisation m designed for large-scale, high-performance simulations; with cells represented as spheres or cylinders with specific interaction force models for each. BioCellion [9] also uses an off-lattice approach, representing cells as spheres or cylinders. Similarly, CellSys [10] implements an off-lattice agent-based framework, modelling individual cells as isotropic, elastic, adhesive spheres.

While the aforementioned tools all have their strengths, they typically make certain assumptions or decisions about the representation of the geometry or simulation approach. This comes at the expense of flexibility and being able to choose the best representation for the specific problem a user is trying to model. To put this control back in the modeller’s hands, Chaste provides multiple plug-and-play geometric representations and modelling approaches within a consistent framework. Chaste is a general-purpose simulation package designed for multi-scale, computationally demanding problems in biology and physiology. It currently supports cell- and tissue-level electrophysiology, discrete tissue modelling, and soft tissue modelling, organised into three modules: cell-based, heart [11] and lung. The cell-based module supports 1D, 2D, and 3D simulations using a variety of modelling approaches, including cellular automata, cellular Potts models, overlapping spheres models, Voronoi tessellation models, and immersed-boundary methods. Chaste provides flexible geometric domains, boundary conditions, and force laws. It includes multiple cell-cycle models and allows users to define mutation states that can affect cell proliferation, migration, and adhesion. Additionally, cell-based Chaste supports reaction-diffusion equations for key nutrients or signalling molecules, whose concentrations can influence these cellular behaviours.

Cell-based Chaste provides a generic multiscale modelling framework for cell-based simulations, encompassing processes from the subcellular to the tissue level. Developed with sound software-engineering principles, Chaste functions as a library of largely independent components [12,13]. Its design allows different force laws, cell-cycle models, and integration methods to be used interchangeably. For example, multiple force laws are available and can be substituted with minimal code modification. Unlike more rigid software packages that only offer a single model or mechanical description, Chaste’s flexible architecture enables direct comparison between alternative simulation approaches [14]. This modular, decoupled structure also facilitates GPU acceleration for individual components without the need to re-engineer the entire codebase. As Chaste is widely used across various research domains, accelerating its cell-based module offers significant benefits to a broad range of studies. For example, Chaste has been used to model intestinal organoids [15], the interplay between mechanochemical patterning and glassy dynamics in cellular monolayers [16], spatial and phenotypic heterogeneity in tumour-macrophage interactions [17], to provide statistical insight into parameterisation of cell-based models of growing tissues [18], cross-talk between Hippo and Wnt signalling pathways in intestinal crypts [19] and cell proliferation and migration during wound healing [20].

As cell-based modelling matures, there is growing demand for simulating larger cell populations and running multiple instances of models that include stochasticity or parameter variation. High-performance computing (HPC) systems aim to meet this need [21]. The increasing reliance on HPC for scientific simulations has driven the development of GPU accelerator technologies and programming models that enable efficient parallel computation. CUDA, developed by NVIDIA, is a low-level platform and API for general-purpose GPU programming that requires explicit memory management and kernel-based execution. OpenCL [22] provides a cross-platform standard for writing portable accelerator code using C-based kernels, offering broad hardware support. Building on OpenCL, SYCL [23] introduces a higher-level C++17-based abstraction for heterogeneous computing, developed by the Khronos Group to improve portability and developer productivity. OpenACC [24] adopts a directive-based approach, allowing developers to offload computations to GPUs with minimal code modifications, similar to OpenMP [25], which also offers directive-based offload to the CPU or GPU.

FLAME GPU 2 [26] is a GPU-accelerated simulator for complex, domain-independent systems. It maps formal agent specifications written in C-based scripting to optimized CUDA code, targeting ABM. FLAME GPU 2 supports multiple agent types, agent communication, and dynamic agent birth and death. In ABM, populations of agents represent individual entities within a simulation, each governed by rule-based behaviours and interactions with other agents and their environment.

Several cell-based simulators have leveraged GPU or HPC technologies for acceleration, often sacrificing extensibility for performance. Biocellion [9] targets massively parallel HPC systems using spatial domain decomposition and MPI to simulate over a billion cells but enforces a fixed one-to-one mapping between cells and agents. BioDynaMo [8] accelerates force calculations via CUDA or OpenCL with pinned memory but supports only a single, non-customisable force law. CellSim3D [27], also CUDA-accelerated, simulates 100,000 cells on a desktop machine but models only mechanical interactions. ya||a [28] employs a spheroid-based CUDA model with spin-like polarities to simulate epithelial sheets and tissue polarity, focusing mainly on morphogenesis. cellGPU [29] accelerates vertex model simulations using CUDA, offering full GPU acceleration for active vertex models and a hybrid GPU/CPU implementation of the self-propelled Voronoi model [30]. ParaCells [31] provides a CUDA-based framework for GPU-oriented cell-centre models with an object-oriented interface that allows users to extend functionality via custom CUDA code. CBMOS [32] is a Python-based, GPU-accelerated cell-centre simulator built on CuPy, while Gell [33] is a hybrid CUDA simulator with a library of predefined cell behaviours.

Previous efforts to accelerate cell-based models in Chaste [34] have primarily focused on distributed computing with MPI [35], mainly to enhance the performance of linear algebra components. While effective, these methods neither exploit massively parallel devices such as GPUs nor accelerate the mechanical aspects of cell-based simulations, and they offer limited flexibility. GPU acceleration is a natural fit for such simulations: large populations of cells exhibit similar behaviours, making the problem well-suited to the GPU computing model. Consequently, GPU acceleration can provide greater performance benefits than traditional parallelisation methods such as MPI. FLAME GPU 2 [26] is a GPU-accelerated simulator for domain-independent complex systems. It maps formal agent specifications written in scripting language to optimized CUDA code for agent-based modelling (ABM). FLAME GPU 2 supports multiple agent types, inter-agent communication, and dynamic agent creation and removal. In ABM, populations of agents represent individual entities within a simulation, each governed by rule-based behaviours and interactions with other agents and their environment.

In this paper, we present a proof of concept for flexible, high-performance, and extensible GPU-accelerated cell-based simulations. This is achieved by integrating GPU acceleration for cell-centre simulations within Chaste using FLAME GPU 2. FLAME GPU 2 was chosen as the GPU acceleration framework for its flexibility and ability to generate well-optimised GPU code without requiring expert-level GPU programming knowledge. This integration provides a separation of concerns: FLAME GPU 2 developers can focus on hardware optimization through a shared agent-based approach, while Chaste developers can concentrate on model development. The framework’s built-in support for spatially partitioned agent-based simulations makes it particularly suitable for off-lattice cell-based models. Its C-like API also reduces the need for specialist GPU expertise among Chaste developers, simplifying the implementation of new GPU-accelerated features and supporting the goal of a flexible, high-performance simulator. Moreover, FLAME GPU 2’s architecture promotes parallel-friendly algorithms through its agent functions and message-passing interface, often achieving performance comparable to hand-written GPU code. Our approach enables the acceleration of multiple models without requiring users to write GPU-specific code. We demonstrate that offloading mechanical computations to the GPU yields substantial speedups and improved mechanical accuracy, all without extensive modification of the pre-existing core Chaste codebase as the GPU-related code is introduced in new classes. The use of FLAME GPU 2 avoids the introduction of HPC-specific code paths and optimisations.

## METHODS

### Reference simulation

2D and 3D reference simulations were constructed to compare the results and performance of the CPU-based simulator with its GPU-accelerated counterpart. These simulations used a well-established ‘cell-centre’ approach [36] in which the location of each cell *i* is given by a single point, **x**_*i*_, representing its centre, and the total force on any cell is a function of the set of cell centres. To determine cell connectivity, we used an ‘overlapping spheres’ approach, whereby two cells interact mechanically if they are within a certain distance of each other. The force **f**_*i*,*j*_ on cell *i* due to cell *j* is assumed to act in the direction of the vector connecting the cells,

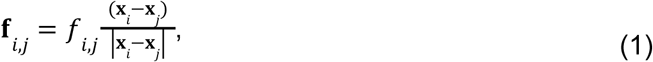

where *f*_*i*,*j*_ is the signed magnitude of **f**_*i*,*j*_. The total force on cell *i* is given by 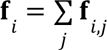, where the sum is over all cells *j* connected to cell *i*. This force is assumed to be balanced by a viscous drag as the cells move, so that

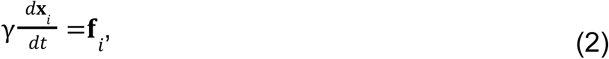

where γ is the drag coefficient. We use a simple forward Euler discretization of Equation (2), so that the position of a cell at time *t* + Δ*t*, given its position at time *t*, is given by

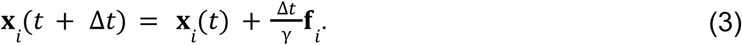

We verified that the time step chosen for all simulations, Δ*t* = 1/12 h, was suitably small for given parameter values to ensure numerical stability.

The signed magnitude of the cell-cell interaction force is given by the function [14]

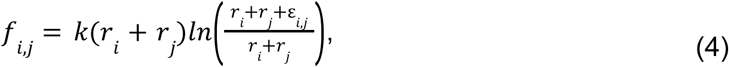

for ε_*i*,*j*_ < 0 and

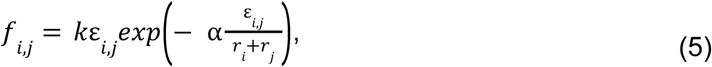

for ε_*i*,*j*_ ≥ 0, where the overlap ε_*i*,*j*_ between the two cells is defined by

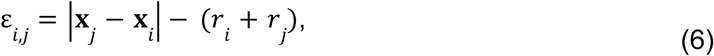

and *r*_*i*_ and *r*_*j*_ are the natural radii of the cells. Hereafter, we assume the following parameter values: γ = 1 α = 5, and *r*_*i*_ = 0. 5 for all *i*.

The simulation began with cells arranged uniformly in a square lattice in 2D or a cubic lattice in 3D with a spacing of 0.66 units. The domain was sufficiently large to prevent cells from reaching its boundaries, effectively rendering it unbounded.

Unless otherwise stated, the reference simulation does not model cell growth or division for the sake of simplicity. Cell division is supported, but is typically disabled to ensure maximum consistency between repetitions of each simulation. No boundary conditions are enforced,

i.e. the cells are free to move anywhere within the domain.

### Targets for GPU acceleration

#### Force resolution

Force resolution is the computation of forces between nodes in node-based simulations. Within cell-based Chaste, this is handled through a set of force classes. The specific force class is selected by the user, however they share a common interface. Each class has an inherited method which accepts as arguments the indices of two nodes and a reference to the cell population containing them. This method is called for each pair of nodes which are close enough to interact. The operation depends only upon the two interacting nodes and doesn’t require the modification or writing of any data to the GPU VRAM during the computation of the force. As such it is well suited to parallelisation. In addition, only nearby nodes are generally interacting, meaning use of spatial data structures for localised interactions also provides significant benefit. A typical individual force computation involves finding the distance between the two nodes. This is used in combination with some material parameters or empirically determined parameters to compute the final force between the two nodes. For any given node, the resultant forces from each of its pairs are summed.

#### Force integration

Force integration is the process of computing how far each node should move based on the resultant forces acting upon it. Cell-based chaste uses a forward euler integration scheme to do this. This simply involves multiplying the resultant force by the timestep and updating the node’s position by the result. There is also within chaste a subsequent check to ensure that no node has moved too far within a single timestep. The computation for each node is completely independent, so this can be parallelised very effectively.

#### Boundary conditions

Boundary conditions are applied after force integration. Some boundary conditions (PlaneBoundaryCondition, SphereGeometryBoundaryCondition) depend only on the boundary condition parameters and individual nodes.

### Integration with FLAME GPU 2

Cell-based Chaste maps to agent-based modelling with cells (and nodes in node-based simulations) being represented by agents in a 2D or 3D environment. Simulation components such as force resolution are handled through the ABM’s behaviours, named *agent functions* within FLAME GPU 2. In FLAME GPU 2, these agent functions are written in C++ using the FLAME GPU 2 API which abstracts the GPU programming model from users. Interaction between agents is indirect via messages with FLAME GPU 2 supporting various message types each. Each message type provides optimisations of data structures to accelerate message querying. I.e. Spatial messaging provides spatial data structures suitable for localised spatial queries.

#### Overview of Chaste simulation flow

A Chaste simulation consists of a series of independently evaluated timesteps. Within each time step, biological processes are simulated, then mechanical interactions are simulated, and finally, *modifiers* are applied. Modifiers can be used to add custom functionality to the simulation and are able to access most elements of the simulation. A typical program flow for a simulation is as follows:

1. AbstractCellBasedSimulation::Solve is the main method
2. Calls `SetupSolve` on each modifier
3. Main loop

a. UpdateCellPopulation

i. removes dead cells
ii. cell division
iii. calls cell population update
b. `UpdateAtEndOfTimestep` for each topology modifier
c. `UpdateCellLocationsAndTopology`

i. defers to numerical method & boundary conditions
d. `UpdateAtEndOfTimestep` for each modifier

#### Use of modifier to contain GPU code

To accelerate the code with minimal modification to the existing Chaste code, the FLAME GPU 2 simulation is embedded within a Chaste simulation modifier. At program start, the FLAME GPU 2 model is constructed. This defines the agents and messages which will be used in the accelerated component. Each cell in the Chaste simulation is represented by a single agent within the FLAME GPU 2 simulation.

FLAME GPU 2 automatically maps the representation of each cell within the modifier to the appropriate CUDA calls at runtime.

The FLAME GPU 2 accelerated, modified program flow is as follows:

1. AbstractCellBasedSimulation::Solve is the main method
2. Calls `SetupSolve` on each modifier **<**- **Create a FLAME GPU 2 Simulation**
3. Main loop

a. UpdateCellPopulation

i. removes dead cells
ii. cell division
iii. calls cell population update
b. `UpdateAtEndOfTimestep` for each topology modifier
c. UpdateCellLocationsAndTopology

i. defers to numerical method & boundary conditions **<**- **Don’t use this**
d. `UpdateAtEndOfTimestep` for each modifier **<**- **Transfer data, perform simulation step with FLAME GPU 2**

At each timestep, *UpdateAtEndOfTimestep* is called on the GPU modifier. The cell data is then retrieved from the Chaste simulation and used to initialise the conditions for the FLAME GPU 2 simulation step. The simulation step itself is then performed, and subsequently the data is retrieved and the Chaste simulation is updated in a similar fashion.

#### Implementation of force calculations

In order to facilitate high performance on the GPU, FLAME GPU 2 uses a message passing interface for communication between agents. In this model, the force calculation maps to two FLAME GPU 2 *agent functions*. In the first function, each agent/cell outputs their location in a message. The second agent function contains the actual logic for the force computation. Each cell will then read the locations of nearby cells from the message list and compute the relevant force contribution. FLAME GPU 2 automatically spatially partitions the domain and only supplies messages from nearby cells which fall within the given interaction radius.

#### Implementation of force integration

Integration in Chaste is handled using a forward Euler integrator. This is implemented in a third FLAME GPU 2 agent function which operates on the cell agents. As the force calculation and integration steps happen sequentially, there is no need to transfer the data back to the CPU host program between these two steps.

### Hardware implementation

All results presented were generated using the University of Sheffield’s *Bessemer* High-Performance Computing cluster [37]. Simulations were conducted on a single GPU node with a single *Intel Xeon Gold 6138 (2.00GHz)* CPU and a single *NVIDIA Tesla V100* GPU allocated.

### Validation

Validation is conducted via a series of tests which compare the CPU and GPU mechanical computations. These are the force calculation and the force integration. The force calculation is tested in both 2D and 3D, for pairs of cells and clusters of cells for a range of cell densities in space in the configurations shown in **Table 1**.

**Table 1.** The configurations for the validation tests. The tests cover two or more cells, in or outside of the interaction radius in both 2D and 3D.

| Test name | Number of cells | Cell interaction radius | Distance between cells |
| --- | --- | --- | --- |
| Two cells in range (2D) | 2 | 1.5 | 0.28 |
| Two cells out of range (2D) | 2 | 1.5 | 2 |
| Multiple cells in range (2D) | 16 | 1.5 | Variable |
| Two cells in range (3D) | 2 | 1.5 | 0.35 |
| Two cells out of range (3D) | 2 | 1.5 | 2 |
| Multiple cells in range (3D) | 64 | 1.5 | Variable |

## RESULTS

### Validation and performance analysis of Chaste

The validation tests set out above in **Table 1** produced no significant errors across any of the test cases (two cells in range, two cells out of range, and multiple cells in range each in 2D and 3D). In each case, there was no significant difference in the results, measured according to the mean difference of the final positions of the cells from those in the reference CPU simulations. In some cases there were minor errors in the order of 10^-8^, likely due to floating point errors.

Figure 1 shows the proportion of runtime spent on mechanics computation in the 2D reference simulation across different cell population sizes. For very small populations, simulations completed rapidly, amplifying relative timing inaccuracies and producing the noise demonstrated in the 1-100 cell range. Beyond this range,mechanics computations consistently account for about 80% of the total runtime. This proportion can vary with the simulation’s initial configuration, but the reference simulation is chosen to represent typical usage. Amdahl’s law predicts the maximum theoretical speedup of a task which is partially parallelised, i.e. it has parallel and serial portions. As the parallel part of the task is further accelerated, returns are diminishing as the serial portion of the task becomes a larger and larger part of the overall runtime of the task. According to Amdahl’s law, if the mechanics portion were perfectly parallelized, an optimal overall speedup of 5x could be achieved.

**Figure 1.**
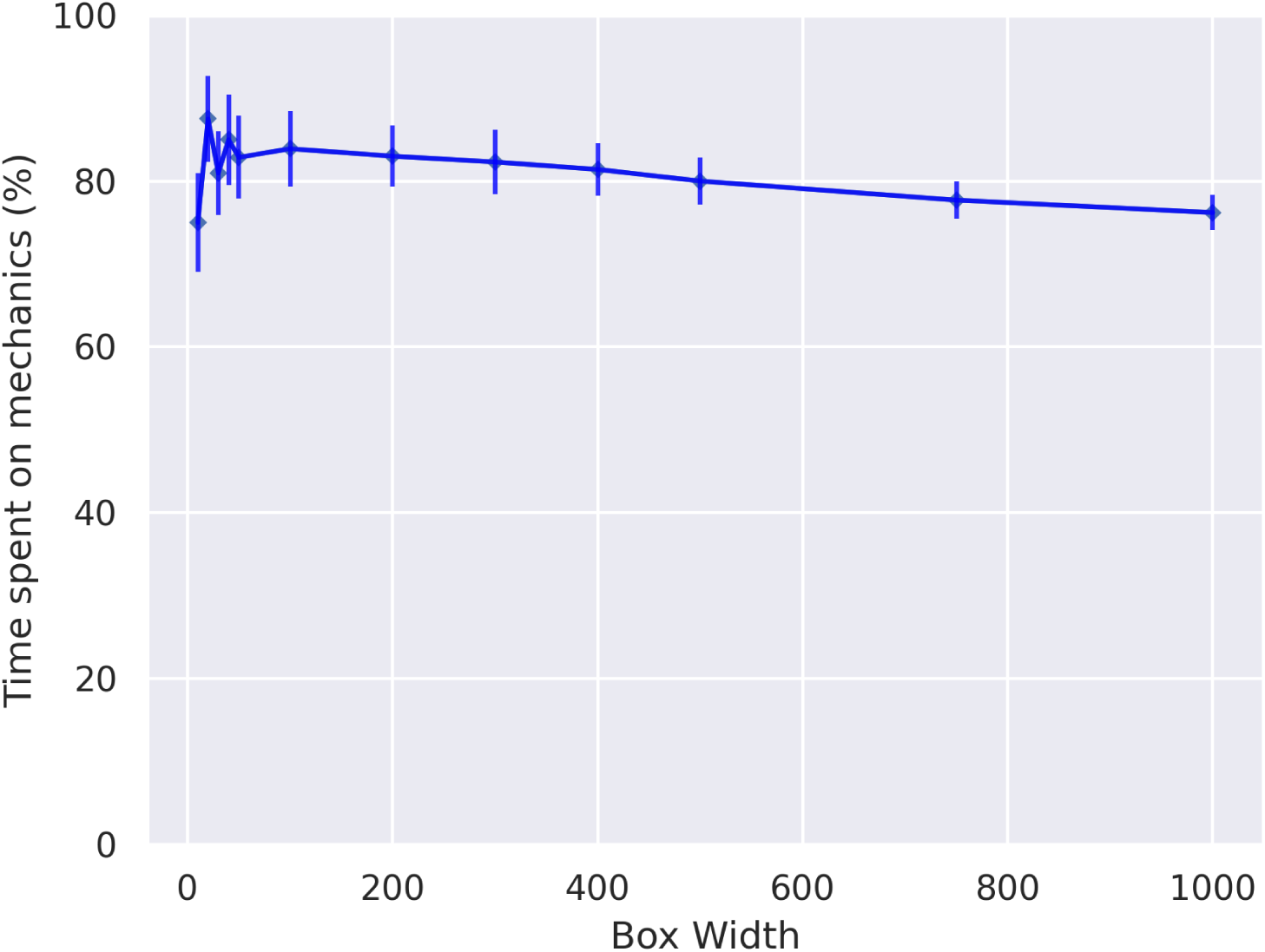
The proportion of runtime spent on mechanics computation in the 3D CPU reference implementations. Error bars show the standard deviation over 10 repetitions of the simulation. This shows the mechanics computation dominates runtime, taking 75% - 85% of the total. This implies a theoretical maximum speedup of 4x-5.5x depending on the specific box size.

### Speedup

Figure 2 shows the actual and theoretical maximum speedups for both 2D and 3D simulations. Theoretical speedups were calculated based on the proportion of time spent on mechanics in the reference simulations, assuming that the mechanics portion of the code is fully parallelizable. In practice, these theoretical maxima are rarely achieved due to hardware or software limitations. In the 2D simulation, the accelerated code achieved a maximum speedup of approximately 2.5x, which is 78% of the theoretical maximum of 3.2x. In the 3D simulation, the maximum speedup was 3.73x, or 90.3% of the theoretical maximum of 4.13x. The overall speedup is also affected by the hardware on which the simulation is run. This will change the relative and absolute performance of the CPU and GPU portions of the code. For example, when tested on a consumer grade NVIDIA RTX 2060, a maximum overall speedup of 4.5x was achieved.

**Figure 2.**
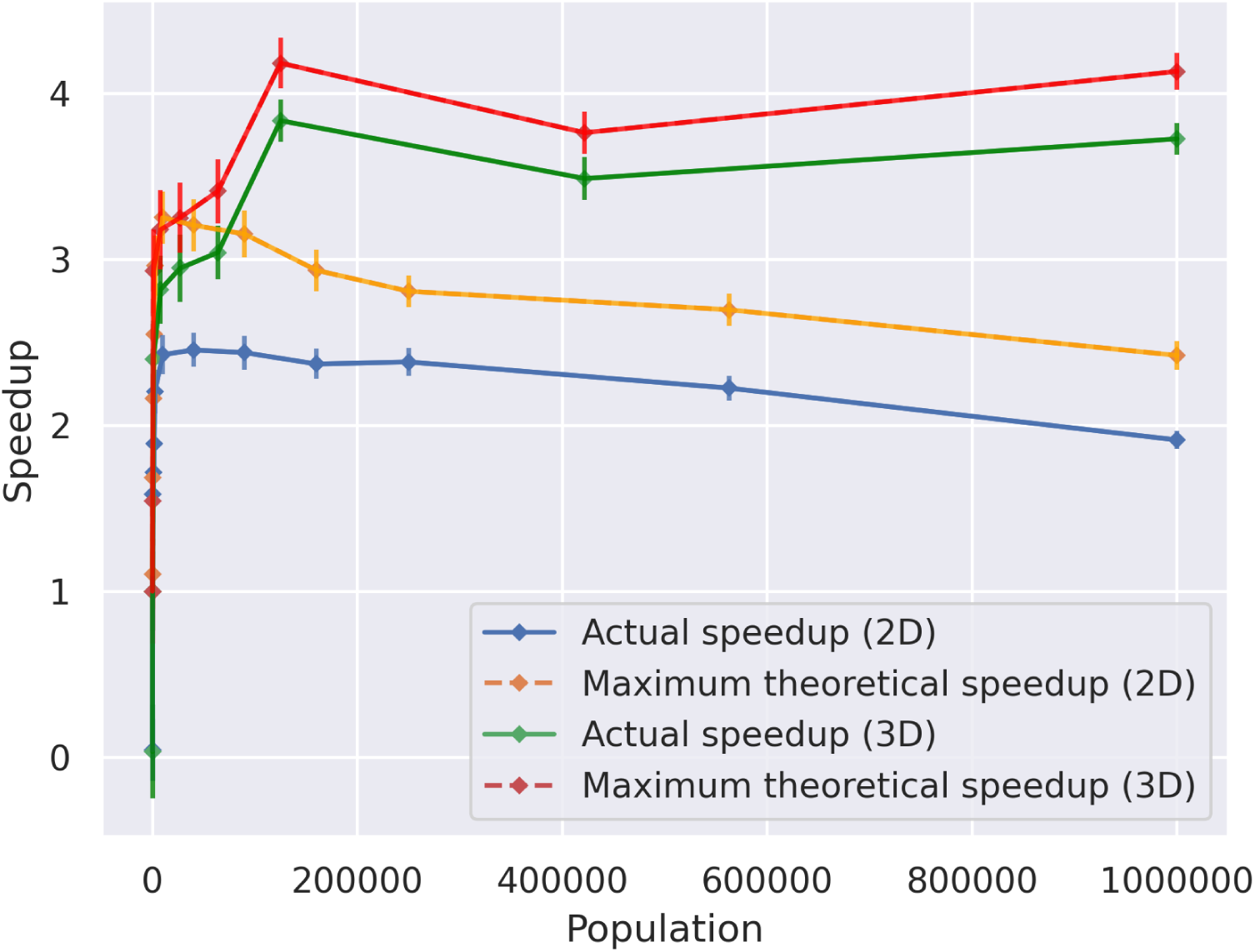
Overall speedup of the 2D and 3D representative simulations. Error bars show the standard deviation over 10 repetitions of the simulation. In 2D, the accelerated version approaches a maximum speedup of around 2.5x, close to the maximum theoretical speedup of ∼3.2x. In 3D, the accelerated version approaches a maximum speedup of around 3.73x, close to the theoretical maximum of ∼4.13x.

The speedup for the force calculation component can be derived from the overall speedup and the fraction of each timestep spent on force computations.

Figure 3 shows the speedups achieved for the mechanics portion of the code. Both 2D and 3D simulations achieved substantial improvements: in 2D, force calculations were accelerated by 24x, while in 3D, the speedup reached 93.6x.

**Figure 3.**
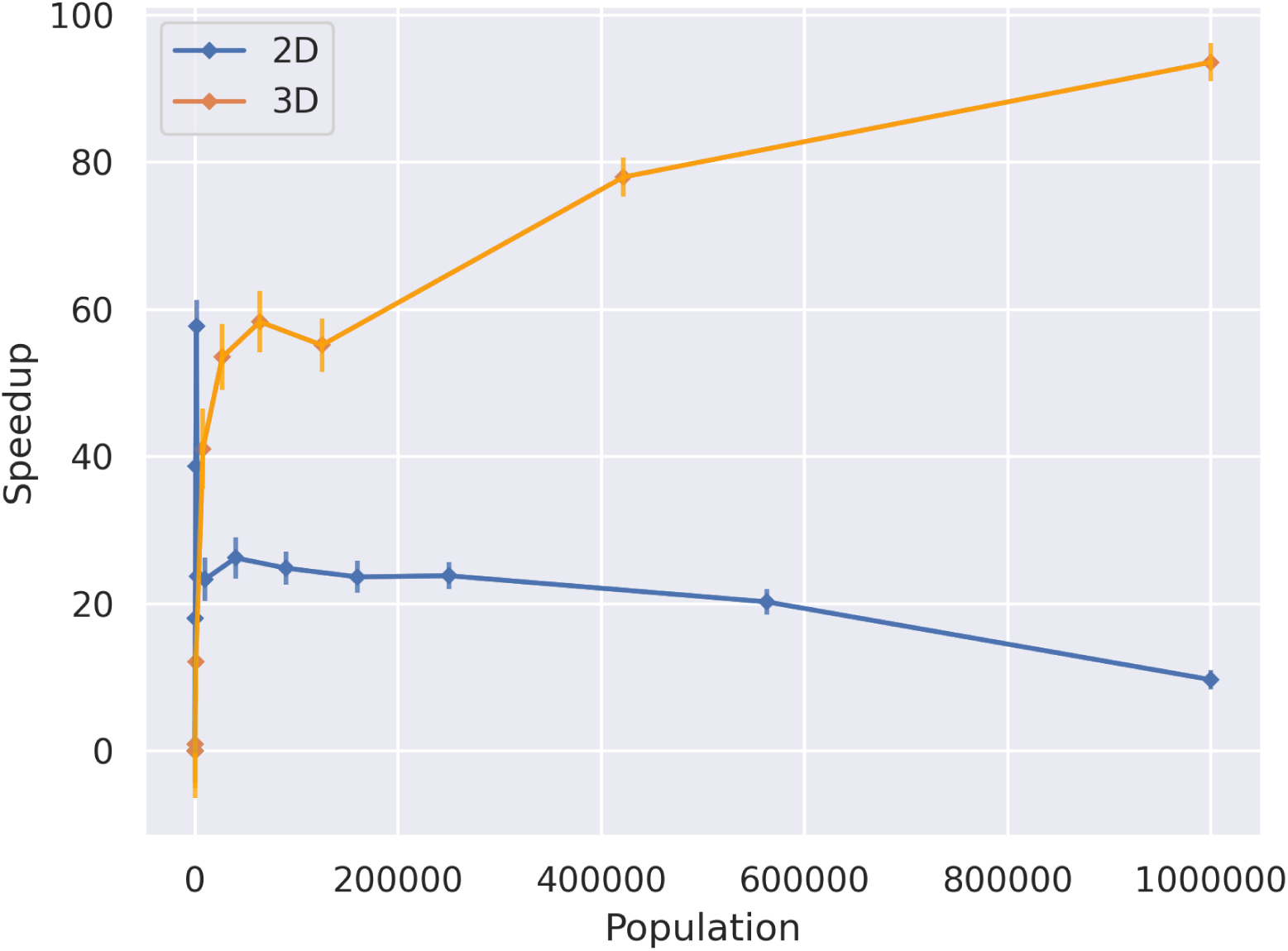
Speedup achieved for the mechanics component of the simulation, i.e. force computation and integration. Error bars show the standard deviation over 10 repetitions of the simulation. In 2D, a speedup of 24x is achieved. In 3D, a speedup of 58x is achieved.

### GPU-accelerated overall profile

Figure 4 breaks down where simulation time is spent in the 3D GPU-accelerated simulation. At very small population sizes, data transfer latency to and from the GPU is significant. For larger populations, CPU time becomes dominant, primarily due to biological updates such as the cell cycle model. For very large populations, the GPU reaches full occupancy, forcing serialisation of kernel executions. At this stage, the compute component dominates runtime.

**Figure 4.**
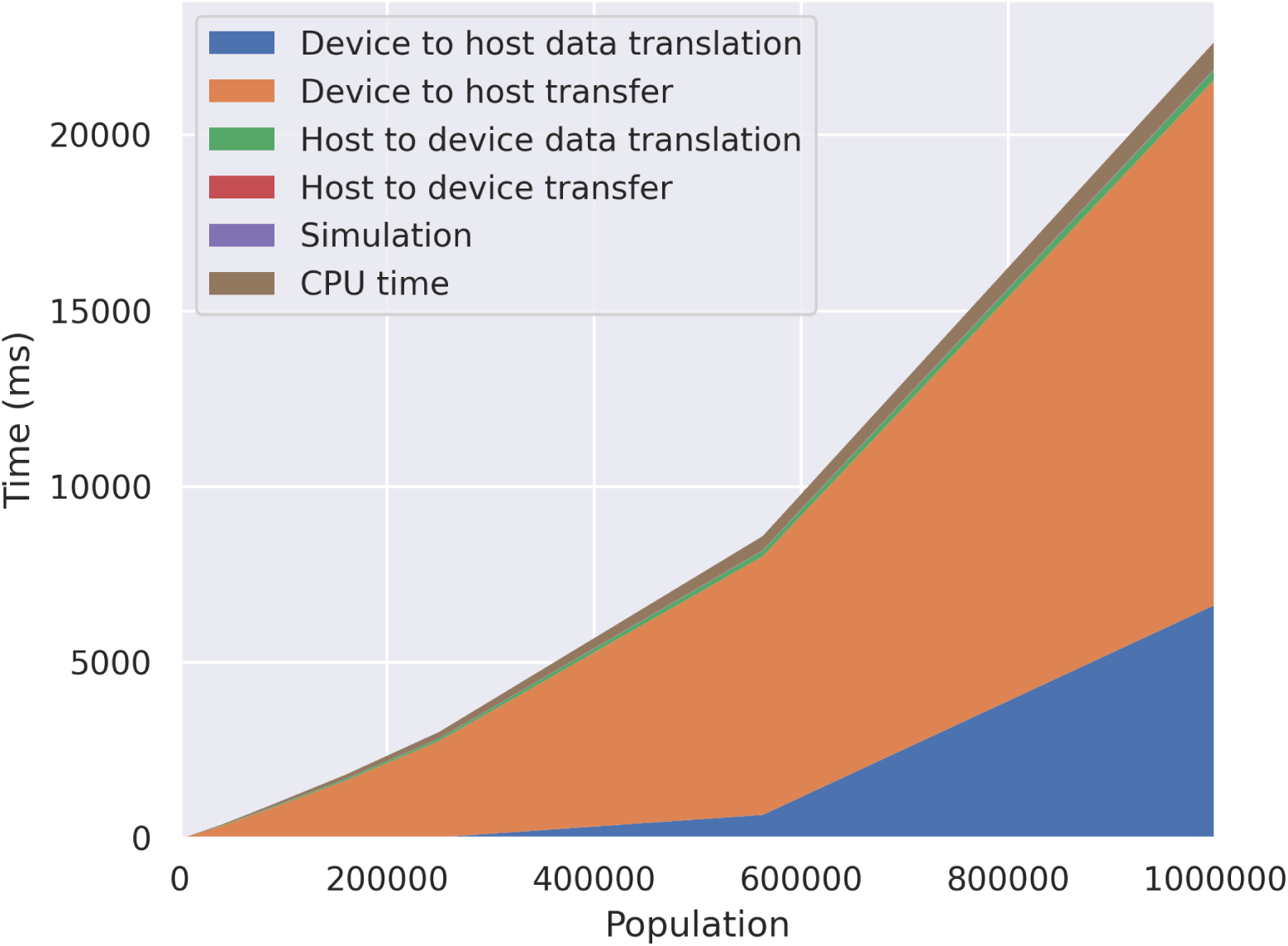
A breakdown of the proportion of time spent on each aspect of the 3D GPU simulation. For very small box widths (<200) the overheads of transferring data to and from the GPU are significant. For larger box widths, the CPU time becomes dominant. At larger population sizes, the compute component dominates the runtime.

Figure 5 shows the proportion of total runtime spent on the CPU for a 2D GPU-accelerated simulation. CPU time accounts for 70–90% of the total runtime, indicating that the mechanics portion of the code has been effectively parallelized. Further significant performance gains would likely require porting more of the code to the GPU.

**Figure 5.**
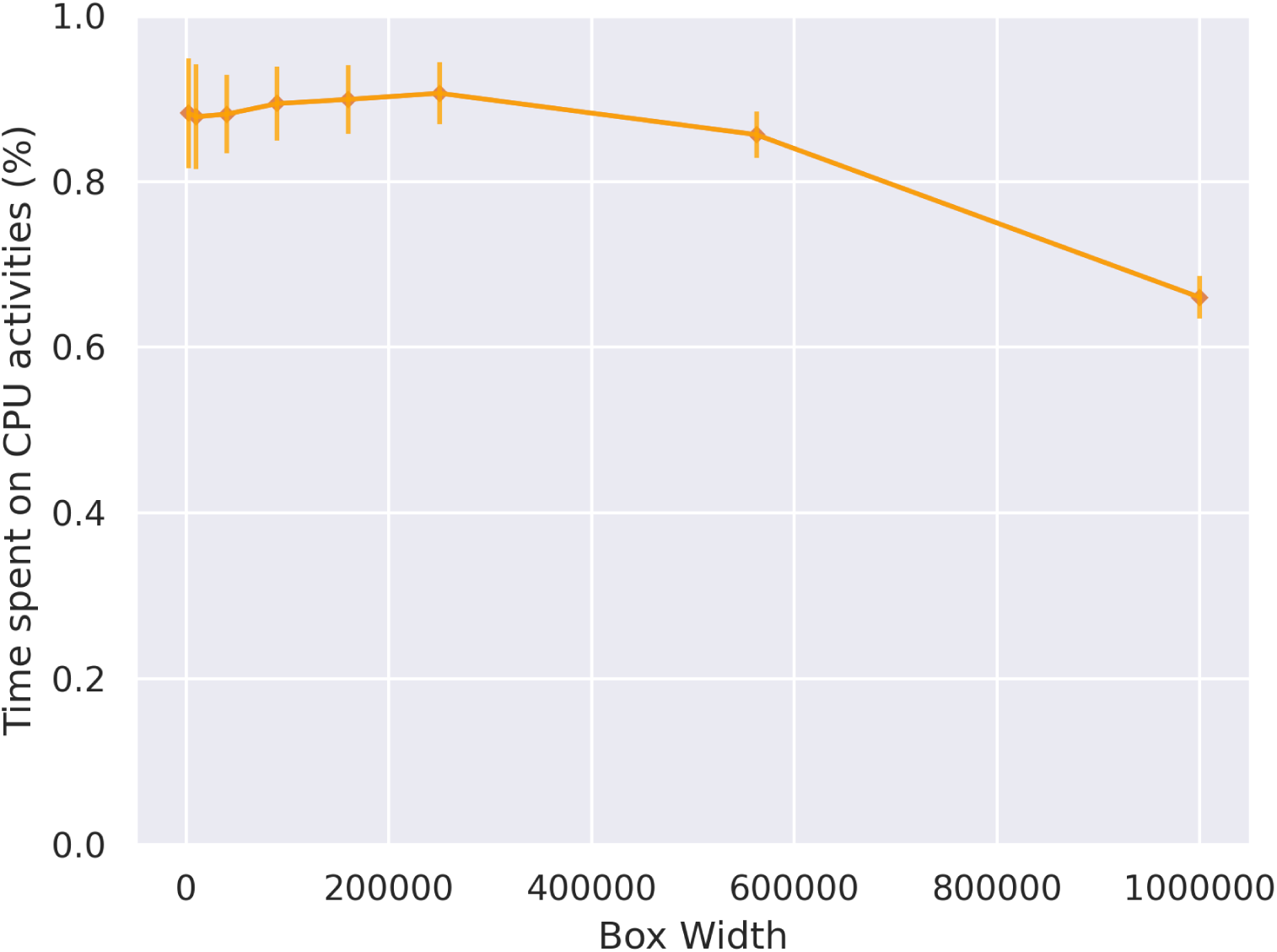
The proportion of total runtime spent by the CPU for the 2D GPU accelerated simulation. Error bars show the standard deviation over 10 repetitions of the simulation. The CPU component is dominating the runtime, indicating that to further significantly improve overall speedup, more code would need to be ported to the GPU.

Figure 6 shows where time is spent within the GPU-accelerated portion of the simulation. For lower population counts, data translation dominates, involving access to cell data via the Chaste API and updating FLAME GPU 2 agents. For larger populations, compute steps become serialized as GPU occupancy reaches its maximum, making the simulation compute-bound. Further performance improvements would therefore require more powerful hardware, algorithmic enhancements, or additional kernel optimization. If Chaste were converted to use a similar memory architecture to FLAME GPU 2, i.e. structure of arrays rather than array of structures, then much of the data translation step could be avoided.

**Figure 6.**
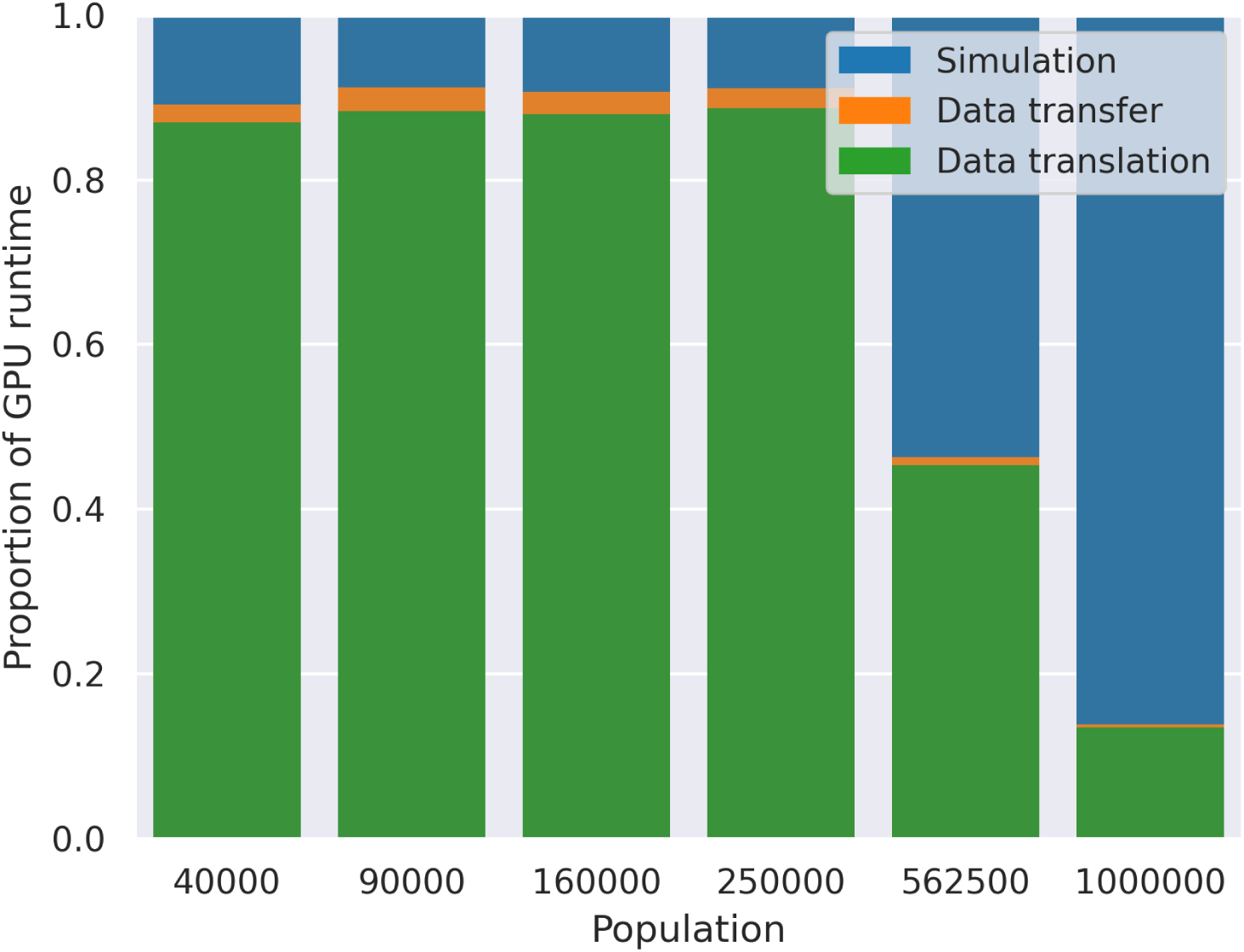
The data transfer, simulation and data translation components of the FLAME GPU 2 part of the simulation for a single 2D simulation. For population sizes which fit within the device comfortably, the data translation dominates the time spent. For larger populations, the compute step becomes dominant.

### Increased mechanical resolution

Since the force computation and simulation step is dominated by data translation, it is possible to increase the number of mechanics timesteps without significantly affecting overall runtime. Multiple force computation and integration steps can be performed without repeatedly transferring data between them.

For a 2D simulation with approximately 10,000 cells, the GPU implementation can complete 300 mechanics timesteps per biological timestep while maintaining runtime parity with the CPU implementation, which performs only a single mechanics step.

Figure 7 shows the percentage error in the position of a cell in a simulation containing two particles with initial x positions of −0.7 and 0.7, and y positions of 0.0. The simulation uses Chaste’s default timestep of 5 minutes and runs until t = 0.5 hours. Performing multiple mechanics timesteps per biological step clearly reduces error, with the maximum position error dropping from 0.00239 to 7.44 × 10^−6^. By the half-hour mark, all trial substeps have converged.

**Figure 7.**
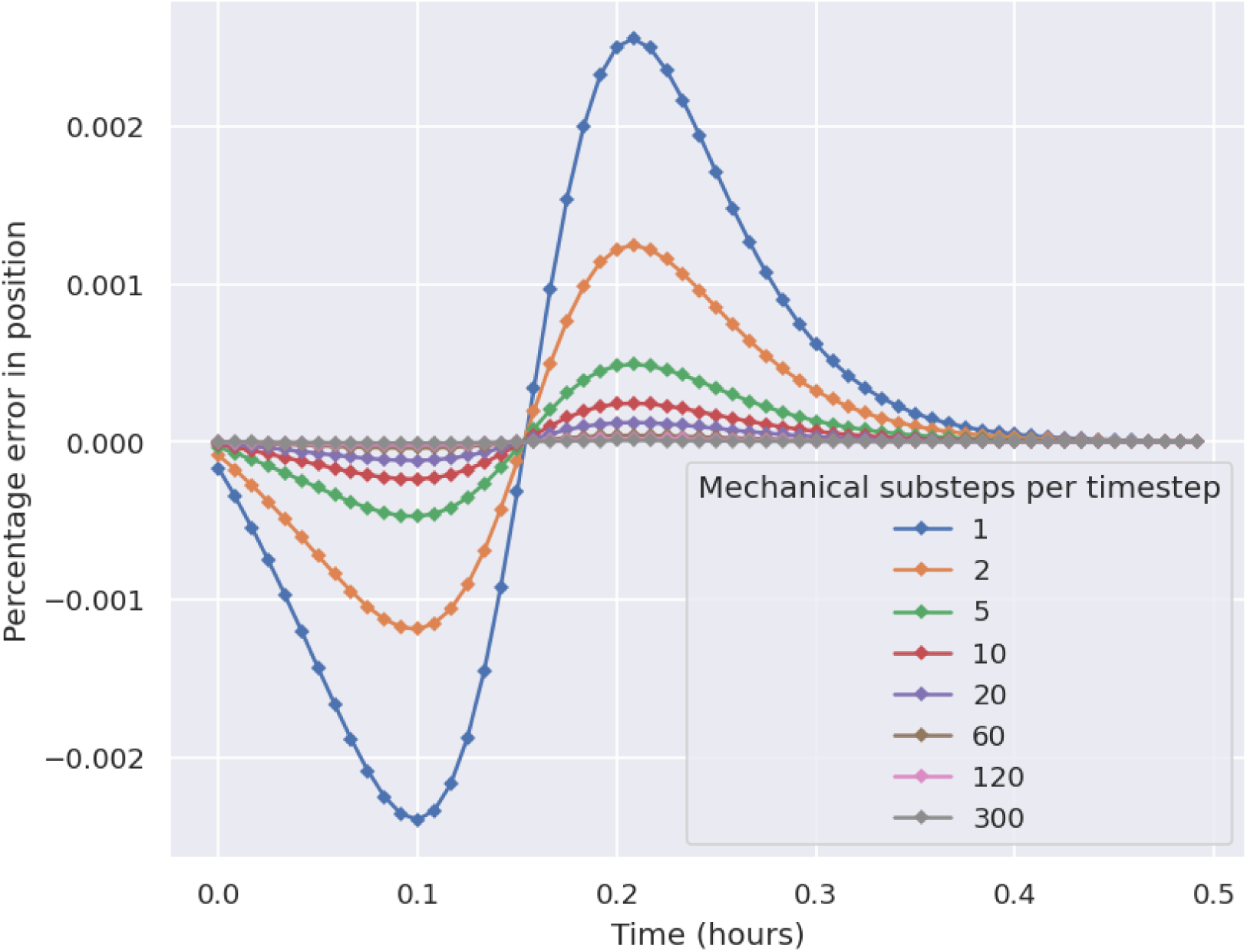
The percentage error in the position of the center of a single cell over time, within a single two cell simulation. Increasing the number of mechanics substeps decreases the percentage error compared to an analytic solution.

This indicates that if only the equilibrium position is of interest, fewer mechanics timesteps may suffice for simulations that converge to equilibrium, provided the total runtime is long enough. However, for simulations focusing on system dynamics or those that do not converge (e.g. with continual cell division), using more mechanics steps can substantially reduce error accumulation. Notably, a simulation using 300 mechanics substeps achieves a runtime roughly equivalent to a CPU simulation using a single mechanics step per biological timestep.

## DISCUSSION

In this paper, we present GPU acceleration for the mechanical component of off-lattice cell-based models in the Chaste library using the agent-based modelling software FLAME GPU 2. We demonstrate that offloading the force computation and integration stages is not only feasible with minimal modifications to Chaste, but also provides significant performance improvements.

Chaste’s simulation modifiers allow end users to leverage GPU acceleration with minimal changes to existing models. Using FLAME GPU 2 reduces the expertise required to implement and maintain new GPU functionality. Tests show no significant loss of precision between GPU-accelerated and CPU versions of the simulations. The performance gains enable simulations of larger cell populations or more repetitions of stochastic simulations in the same duration, increasing confidence in the results. Running more simulations within the same timeframe also facilitates parameter exploration and fitting, which is often critical for model evaluation. Additionally, performing multiple mechanics timesteps can improve simulation accuracy without increasing runtime, particularly for systems where dynamics are studied or equilibrium is not reached.

While current performance gains are substantial, they are limited by the remaining serial portions of the code. Future work could focus on parallelising these sections. Moving some or all biological simulation components to the GPU could further improve performance by reducing costly data transfers between CPU and GPU. Moreover, expanding GPU support to a wider range of mechanical models - beyond the current single force model and integrator used as a proof of concept - would make Chaste highly performant and flexible, requiring no GPU programming knowledge from end users. Collecting performance profiles across diverse models and simulation conditions would provide a clearer picture of real-world applicability and potential speedups. Many cell-based simulations also incorporate partial differential equations representing substance availability or concentration, which can be computationally expensive. Accelerating Chaste’s partial differential equation solvers would complement the current GPU work and further enhance performance.

## KEY POINTS

- We implemented GPU acceleration for cell-based simulations in Chaste using FLAME GPU 2, without requiring extensive specialist GPU programming knowledge.
- The acceleration achieved a 92.6x speedup for the targeted components and a 3.73x overall speedup.
- With the accelerated force calculations, multiple physics timesteps can be resolved per biological timestep, significantly reducing integration errors during the model’s mechanical simulations.

## ACKNOWLEDGMENTS

The authors thank Kwabena Amponsah, Fergus Cooper, Jack Jennings, Gary Mirams, and Joe Pitt-Francis for useful discussions about Chaste.

## DATA AVAILABILITY

No external datasets were generated or analysed during the current study. The code necessary to reproduce all the figures presented in the manuscript is available from GitHub at https://github.com/Chaste/gpu-benchmark-2026. Running the code will require a valid chaste installation. Instructions for installing chaste can be found on the website https://chaste.github.io/docs/.

## AUTHOR CONTRIBUTIONS

M.L. developed the new functionality. A.F. and P.R. supervised the project. M.L., A.F., and P.R. wrote and reviewed the manuscript.

